# Radiation-induced cell cycle perturbations: a computational tool validated with flow-cytometry data

**DOI:** 10.1101/2020.07.29.226480

**Authors:** Leonardo Lonati, Sofia Barbieri, Isabella Guardamagna, Andrea Ottolenghi, Giorgio Baiocco

## Abstract

Cell cycle progression can be studied with computational models that allow to describe and predict its perturbation by agents as ionizing radiation or drugs. Such models can then be integrated in tools for pre-clinical/clinical use, *e.g.* to optimize kinetically-based administration protocols of radiation therapy and chemotherapy.

We present a deterministic compartmental model, specifically reproducing how cells that survive radiation exposure are distributed in the cell cycle as a function of dose and time after exposure. Model compartments represent the four cell-cycle phases, as a fuction of DNA content and time. A system of differential equations, whose parameters represent transition rates, division rate and DNA synthesis rate, describes the temporal evolution. Initial model inputs are data from unexposed cells in exponential growth. Perturbation is implemented as an alteration of model parameters that allows to best reproduce cell-cycle profiles post-irradiation. The model is validated with dedicated *in vitro* measurements on human lung fibroblasts (IMR90). Cells were irradiated with 2 and 5 Gy with a Varian 6 MV Clinac at IRCCS Maugeri. Flow cytometry analysis was performed at the RadBioPhys Laboratory (University of Pavia), obtaining cell percentages in each of the four phases in all studied conditions up to 72 hours post-irradiation.

Cells show early G_2_-phase block (increasing in duration as dose increases) and later G_1_-phase accumulation. For each condition, we identified the best sets of model parameters that lead to a good agreement between model and experimental data, varying transition rates from G_1_- to S- and from G_2_- to M-phase.

This work offers a proof-of-concept validation of the new computational tool, opening to its future development and, in perspective, to its integration in a wider framework for clinical use.

**Author summary:** We implemented a computational model able to describe how the progression in the cell cycle is perturbed when cells are exposed to ionizing radiation. It is known that radiation causes delays or arrest in cell cycle progression, and also that cells that are in different phases of the cycle at the time of exposure show different sensitivity to radiation. Chemotherapeutic drugs also affect cell cycle, and their action can be phase-specific. These findings can be exploited to find the optimal protocol of a combined radiotherapy/chemotherapy cancer treatment: to this aim, we need to know not only the effectiveness of an agent (dose/drug) in terms of cell killing, but also how surviving cells are distributed in the cell cycle. With the model we present, this information can be reproduced as a function of dose and time after radiation exposure. To test the model performance we measured distributions of cells in different phases of the cycle (using flow-cytometry) for human healthy fibroblast cells exposed to X-rays. The results of this work constitute a first step for further development of our model and its future integration in a tool for pre-clinical/clinical use.

## Introduction

Cell cycle is a process of fundamental importance, at the basis of cell growth and replication. In eukaryotic cells it is typically divided in four phases: G_1_-phase, in which the cell gets ready to DNA synthesis; S-phase, when DNA replication takes place; G_2_-phase, when the cell begins preparation for mitosis; and M-phase, where cell division takes place. Each phase is characterized by a different length. Various proteins, cyclins and cyclin-dependent kinases, regulate transitions between two phases and block progression in specific checkpoints if the system detects errors [1].

Exposure to ionizing radiation is known to affect cell cycle progression: radiation causes DNA damage, and an arrest in cell cycle progression originates as the cell activates DNA repair mechanisms. A “successful” repair can lead to further progression in the cycle (recovery of the arrest, with a resulting delay), while, if the repair is unsuccessful, the cell can eventually die thus exiting the cycle or progress fixing alteration (genome instability). Cell cycle arrests following radiation exposure have been observed experimentally mainly as accumulation in G_2_/M-phase, followed by possible G_1_-phase arrest. It is also well known that cells in different phases of the cycle at the time of exposure show a different sensitivity to the radiation insult, with cells in S-phase being generally more radioresistant than cells in G_1_ and G_2_, and cells in M being highly radiosensitive. Quiescent cells in a G_0_ state are also radioresistant, as they cannot undergo clonogenic death until brought to re-enter the replicative cycle.

Findings on the interplay between radiation action and cell cycle progression are applied in radiation therapy for cancer treatment. In particular, when a fractionation scheme is used, the total dose is split in smaller fractions: this allows the redistribution of surviving cancer cells within the cell cycle, the repair of sub-lethal damage, the re-oxygenation of the tumor and repopulation of normal and malignant tissues [2]. Chemotherapeutic drugs also affect cell cycle progression, and their action can as well be phase-specific, *e.g.* interfering with replication in the S-phase or damaging the formation or dissociation of the mitotic spindle in the M phase. The treatment effectiveness will be finally dependent on several factors, as the spatial distribution of the tumor cell mass (oxygenation heterogeneity), the timing of the drug/radiation dose delivery, the time between doses, the specific radiosensitivity of the tumor, etc. From the combination of treatment-dependent perturbations of cell-cycle progression and cell-cycle-dependent therapeutic sensitivity we get the rationale behind the use of kinetically-based administration protocols of chemotherapy and radiation therapy: as a general consideration, favouring synchrony and arrest of cells at a particular cell-cycle phase can improve the effectiveness of the next dose of radiation/chemotherapy, administered within an appropriate time so that synchrony/arrest is not lost [3].

For radiation treatments, the modelling of the perturbation of the cell cycle might then be used as an input to refine the evaluation of the Tumour Control Probability (TCP), defined as the probability that no cancer cells clonogenically survive, and to optimize the fractionated treatment protocol in terms of fraction numbers, dose per fraction and time between fractions [4]. Possible synergistic effects of concurrent treatments with radiation and chemotherapeutic drugs have to be explored, especially, in perspective, going from conventional radiotherapy to particle therapy for radioresistant tumours, in both target- and healthy tissues. Clinically driven mathematical models can be used for this purpose as tools to understand, study, and provide useful predictions related to the outcome of various treatment protocols used to treat human malignancies. The use of such tools could speed up delivery of efficacious treatments to patients, providing indications prior to beginning actual testing and long and costly clinical trials, and also preventing the use of potentially sub-optimal treatment combinations [5].

Tools of this kind, to be used in a pre-clinical/clinical framework, have to rely on solid computational models able to describe cell cycle progression and predict the outcome of a given perturbation. Different cell-cycle models have been developed, greatly varying in complexity, from compartmental models based on ordinary differential equations (ODEs), to multi-scale models predicting population growth, possibly taking into account intracellular biochemical processes or factors of the cell environment that affect the fate of each individual cell. Generally speaking, models limited to the prediction of the distribution of cells in the cycle have a deterministic nature, their output being fully determined by parameter values and initial conditions. Different options are available: the model can include explicit expressions for the concentration of regulators of cell cycle progression and their time evolution (usually limited to essential interactions), thus providing a molecular insight on the system [6]. In this case, model parameters are activation and degradation rates of regulatory proteins and their concentration. Alternatively, the model can include expressions for the percentage of cells that are found to be in a given cell cycle phase, thus providing a “population overview” [7]–[8]. Model parameters are then transition probabilities between different phases. The perturbation of the system is finally described by a variation of the values of parameters that govern its evolution. Radiation action can be described as leading to an outcome subject to probability laws, as it is the case for clonogenic cell survival. This would suggest the use of a stochastic model, where the same set of input parameters and initial conditions will lead to an ensemble of different outputs. When describing cell survival coupled to the perturbation of the cell cycle, hybrid models can be implemented, incorporating randomness in the deterministic evolution of the system [5]. The choice of which model and how to implement it necessarily depends on the needs and aims of the study, and above all, on the availability of data for model benchmark.

The purpose of this work is to set the basis of a computational model to specifically describe how cells that survive radiation exposure are distributed in the cell cycle. Flow cytometry is adopted to provide dedicated experimental data for the testing and benchmark of this model. Data are obtained from measurements on healthy human fibroblasts (IMR90 cell line) exposed to X-ray doses up to 5 Gy and followed from 6 hours up to 72 hours post irradiation. Using flow cytometry, the position of the cell in the cycle is primarily determined measuring its DNA content. We therefore present a new implementation from a previously published deterministic model (up to now applied to cell-cycle perturbation by chemotherapeutic drugs only), explicitly taking into account DNA content as a variable, besides the time variable for the temporal evolution of the system [9]. The developed model is able to describe how radiation modifies in time the percentage of cells in all phases of the cycle, with a benchmark also on the distinction of G_2_- and M-phases (at difference with previous implementations). Radiation-induced perturbations and their evolution in time are reproduced varying two of the model parameters, representing transition rates from G_1_- to S-phase and from G_2_- to M-phase. Upon further developments, that we later discuss in great detail, this model can be integrated in the framework of tools to be developed for predictions/optimization with cancer cells and future prediction about cancer therapy, considering the dynamics of cell cycle progression and its alteration by chemotherapeutic drugs and radiation and, eventually, cell death leading to the control of tumour cell population.

## Results

In what follows, we first present experimental measurements obtained with the IMR90 cell line. We then introduce the theoretical model and discuss how model parameters are estimated reproducing the unperturbed experimental condition (cells in exponential growth in absence of radiation exposure). We finally show that modification of such parameteres allows to reproduce cell-cycle perturbation induced by radiation. All details on cell culture and experimental procedures and on the mathematical model and its MATLAB implementation are given in the Materials and Methods section.

### Experimental results

We present experimental data obtained to characterize the human fibroblasts cell line IMR90 in terms of: i) the growth of the cell population as a function of time in the control (unirradiated, sham) condition; ii) how the distribution of vital cells in the different cell-cycle phases is affected, again as a function of time, when cells are either unirradiated or exposed to X-ray doses of 2 and 5 Gy. In all experimental conditions cells are followed in time until 72 hours.

#### Estimation of population doubling time in unirradiated conditions

Table 1 reports the average number of cells in unirradiated conditions for each time point (counted with a Bürker chamber). The number of cells as a function of time is fitted to an exponential function in Fig. 1. Such function allows to consider growth of a cell population to derive an estimation of the doubling time (T_D_), obtained from the fit using the relation:

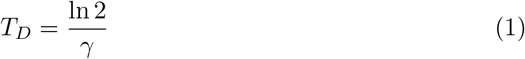

where *γ* is the exponent coefficient of the fitted function (see Materials and Methods). The estimated T_D_ is of 38.4 (CL at 95% [31.7, 48.8]).

**Fig 1.**
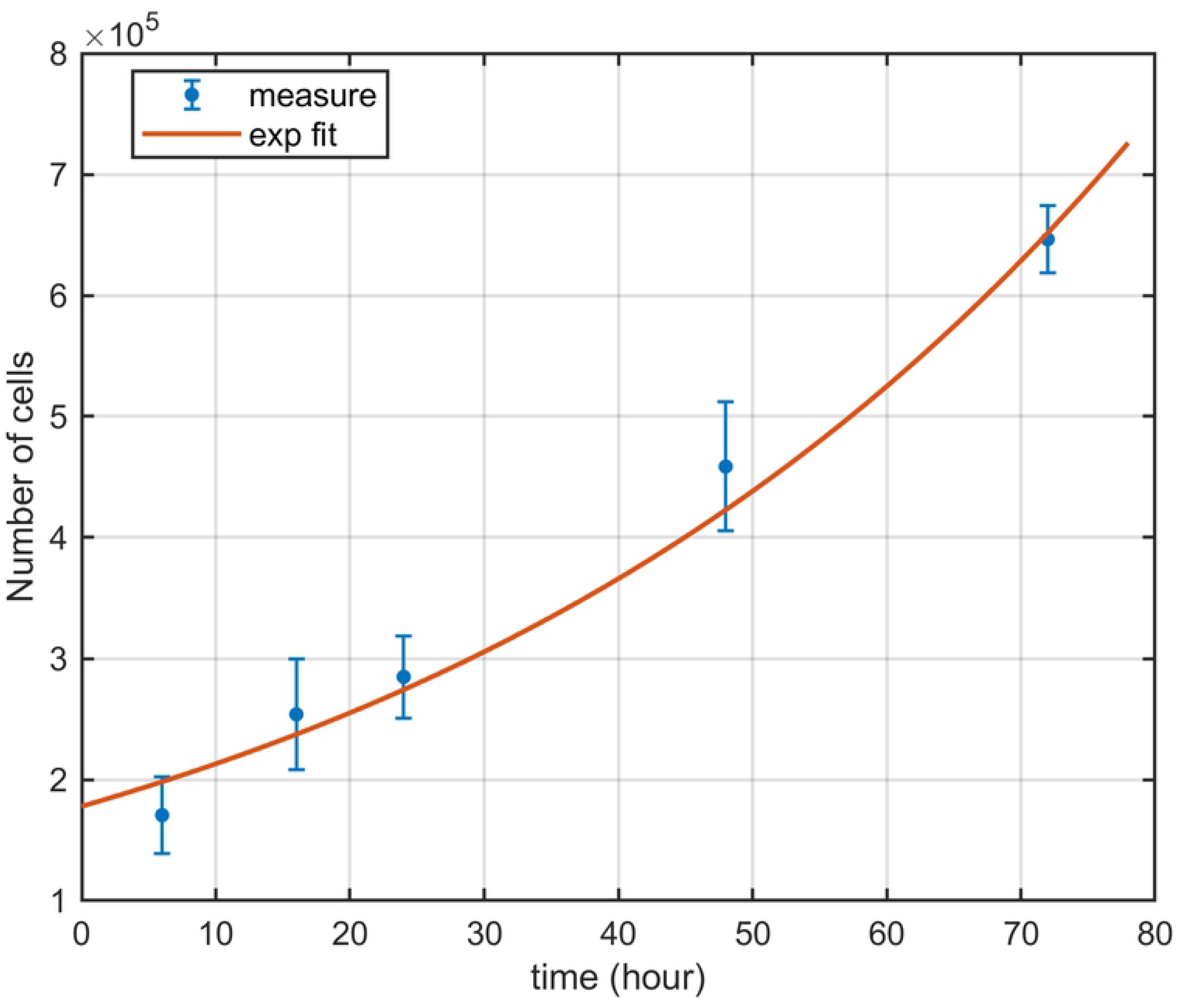
Cell growth curve. Number of unirradiated cells as a function of time and exponential fit of the data.

**Table 1.**
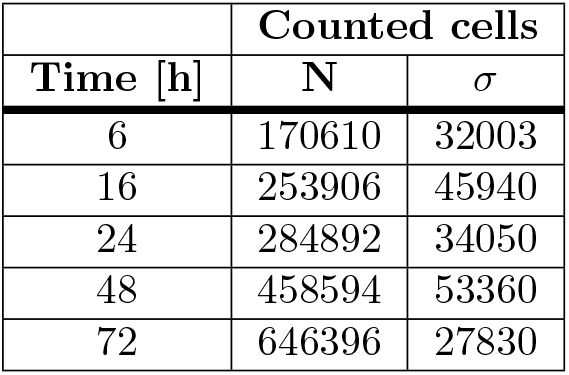
Average number of unirradiated cells counted at each time point with the corresponding standard deviation among replicates.

Data are given as average between both biological (3) and technical (2) replicates with the corresponding standard deviation (1*σ*).

#### Flow-cytometry analysis for cell-cycle distribution

Results on experimental percentages of cells in each phase of the cell cycle for sham, 2 Gy and 5 Gy are given in Table 2. Data are given as average between both biological (3) and technical (2) replicates with the corresponding standard deviation (1*σ*).

**Table 2.** Experimental percentages of cells in each cell cycle phase for different doses (sham, 2 Gy, 5 Gy) at each time point.

| Time<br>[h] | Dose<br>[Gy] | Percentage of cells in: |  |  |  |
| --- | --- | --- | --- | --- | --- |
|  |  | G<sub>1</sub> | S | G<sub>2</sub> | M |
| 0 | 0 | 58.26 ± 1.64 | 31.25 ± 0.48 | 9.35 ± 1.08 | 1.132 ± 0.432 |
| 6 | 0 | 55.30 ± 0.95 | 37.08 ± 1.42 | 7.04 ± 0.81 | 0.582 ± 0.183 |
|  | 2 | 45.78 ± 0.96 | 31.56 ± 1.38 | 22.47 ± 1.60 | 0.189 ± 0.076 |
|  | 5 | 45.63 ± 1.89 | 30.51 ± 2.29 | 23.84 ± 2.81 | 0.018 ± 0.008 |
| 16 | 0 | 62.62 ± 1.47 | 27.38 ± 1.56 | 9.12 ± 1.13 | 0.883 ± 0.226 |
|  | 2 | 82.37 ± 1.43 | 2.32 ± 0.33 | 14.91 ± 1.68 | 0.377 ± 0.118 |
|  | 5 | 60.82 ± 3.15 | 1.23 ± 0.34 | 37.54 ± 3.49 | 0.409 ± 0.054 |
| 24 | 0 | 68.05 ± 2.09 | 24.57 ± 2.12 | 6.77 ± 0.99 | 0.600 ± 0.184 |
|  | 2 | 88.23 ± 1.68 | 2.78 ± 0.37 | 8.91 ± 2.03 | 0.077 ± 0.021 |
|  | 5 | 70.50 ± 3.30 | 1.17 ± 0.43 | 28.26 ± 3.42 | 0.062 ± 0.021 |
| 48 | 0 | 79.65 ± 0.69 | 14.68 ± 0.62 | 5.35 ± 0.60 | 0.319 ± 0.150 |
|  | 2 | 87.49 ± 0.71 | 5.17 ± 1.07 | 7.25 ± 1.75 | 0.084 ± 0.044 |
|  | 5 | 82.75 ± 3.96 | 1.79 ± 0.54 | 15.42 ± 4.50 | 0.031 ± 0.010 |
| 72 | 0 | 79.02 ± 0.46 | 16.84 ± 0.40 | 3.86 ± 0.54 | 0.279 ± 0.086 |
|  | 2 | 82.52 ± 0.67 | 9.16 ± 1.97 | 8.15 ± 1.94 | 0.171 ± 0.059 |
|  | 5 | 76.80 ± 3.77 | 3.20 ± 1.47 | 19.95 ± 4.72 | 0.051 ± 0.023 |

#### Cell cycle distribution in the control condition

For the sham condition we observe in Fig.2 almost constant percentages in cell cycle phases vs. time in the early time points, while at 48 h and 72 h the percentage of cells in G_1_-phase increases from 61% (average value between 0 h and 24 h) to 79%. Data from Fig.2 suggest that already at 48 h cells are approaching confluence, and their distribution in the cell cycle is affected. Based on these results, time points until 24 h only can be considered to have cells in an exponential growth phase.

**Fig 2.**
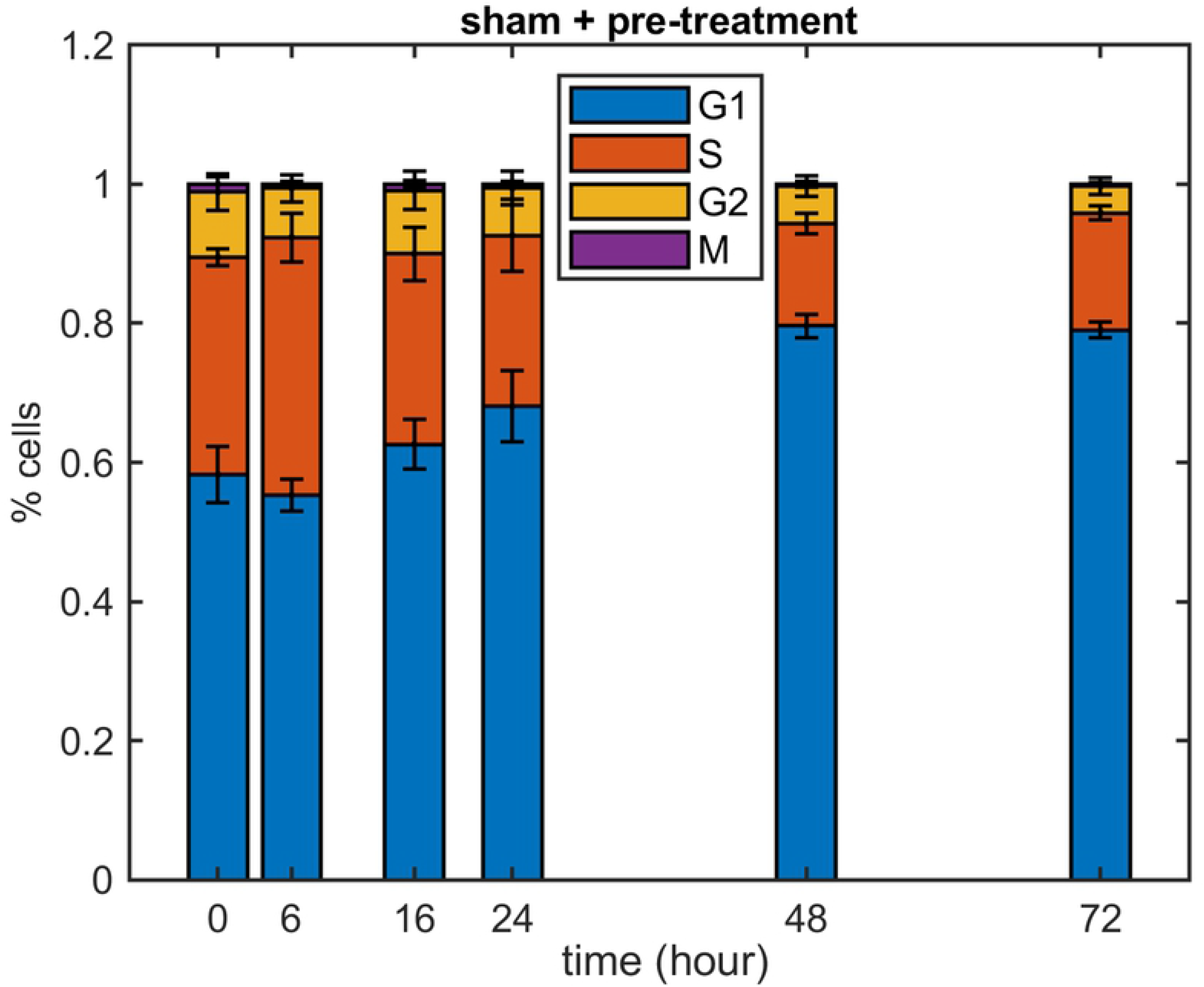
Experimental cell cycle distribution in sham samples. Percentages of cells in the four cell-cycle phases as a function of time for the sham and the pre-treatment condition (data from Table 2). Percentages of cells in the M-phase are generally too low to be appreciated in the plot.

#### Cell cycle distribution after irradiation

Fig.3 shows the percentages of cells in different cell-cycle phases for samples irradiated with 2 Gy and 5 Gy, respectively. Compared to the sham condition, for both doses there is a strong increase of cells in G_2_-phase at the early 6 h time point. A block in G_2_ following radiation exposure is to be attributed to the activation of cell repair mechanisms.

**Fig 3.**
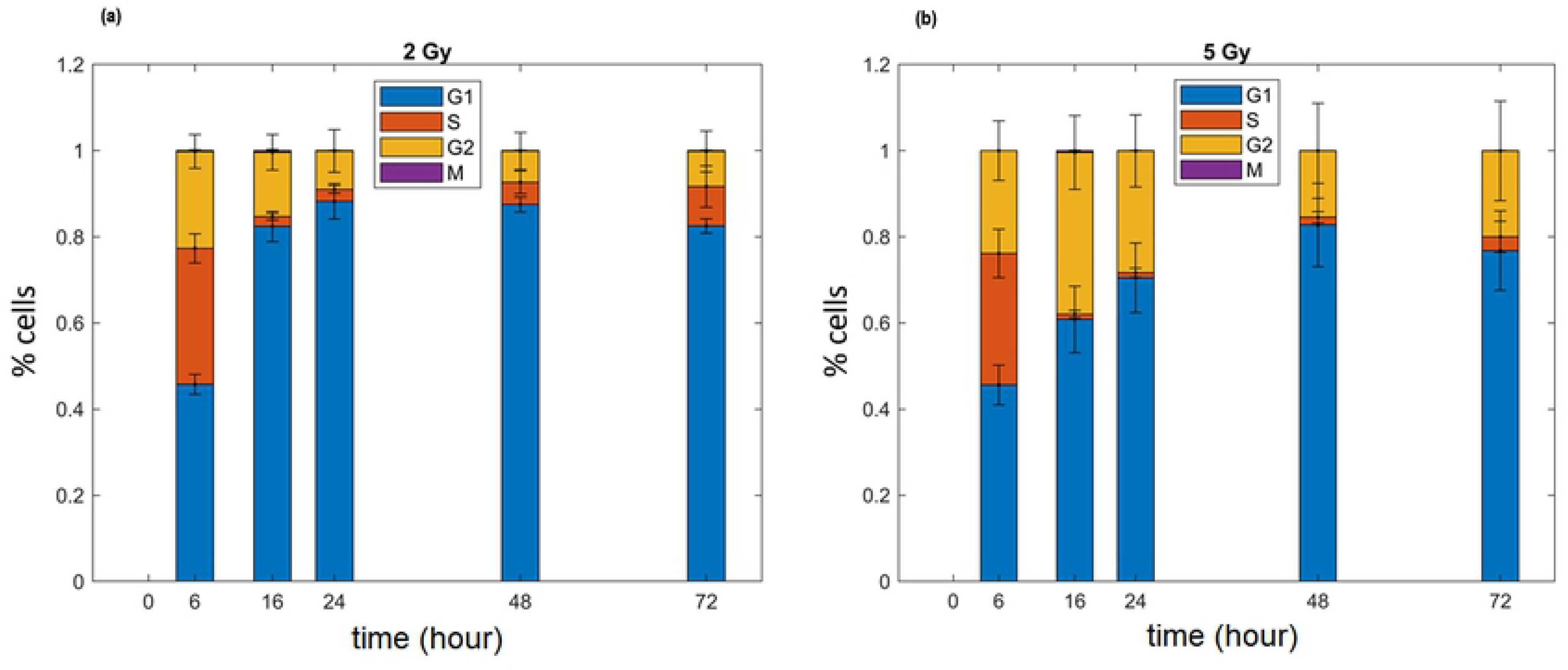
Experimental cell cycle distribution for irradiated samples. Percentages of cells in the four cell-cycle phases as a function of time for the 2 Gy irradiated condition **(a)**, and for the 5 Gy irradiated condition **(b)** (data from Table 2). Percentages of cells in the M-phase are generally too low to be appreciated in the plot.

For the 2 Gy condition, the percentage of cells in G_2_ reaches its maximum at this time point, and later decreases to an average value of 8.10% (between 24 h and 72 h), comparable with the average value for the sham between 0 and 24 h (8.07%).

Correspondingly, the percentage of G_1_-phase cells goes up in the same time-frame to an average value of 85.90%, and the percentage of S-phase is drastically reduced under 10% for any time point.

In the 5 Gy condition, the percentage of cells in G_2_ is maintained higher at all the time points, being maximal at 16 h with a value of 37.54%. A similar increase of G_1_-phase is shown at later time points, with a peak at 48 h. This latter effect can be both attributed to a block in G_1_, and to influx and accumulation in G_1_ of cells surviving the earlier block in G_2_ that return cycling.

#### Control vs. irradiated conditions

In Fig. 4 we report the relative differences in percentages of cells in cell-cycle phases between the sham and the two irradiated conditions. Plotted values are calculated for each time point as:

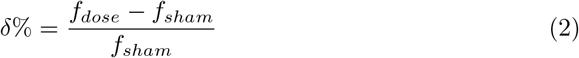

where *f*_*dose*_ and *f*_*sham*_ are, respectively, the percentages for the irradiated condition and for the sham.

**Fig 4.**
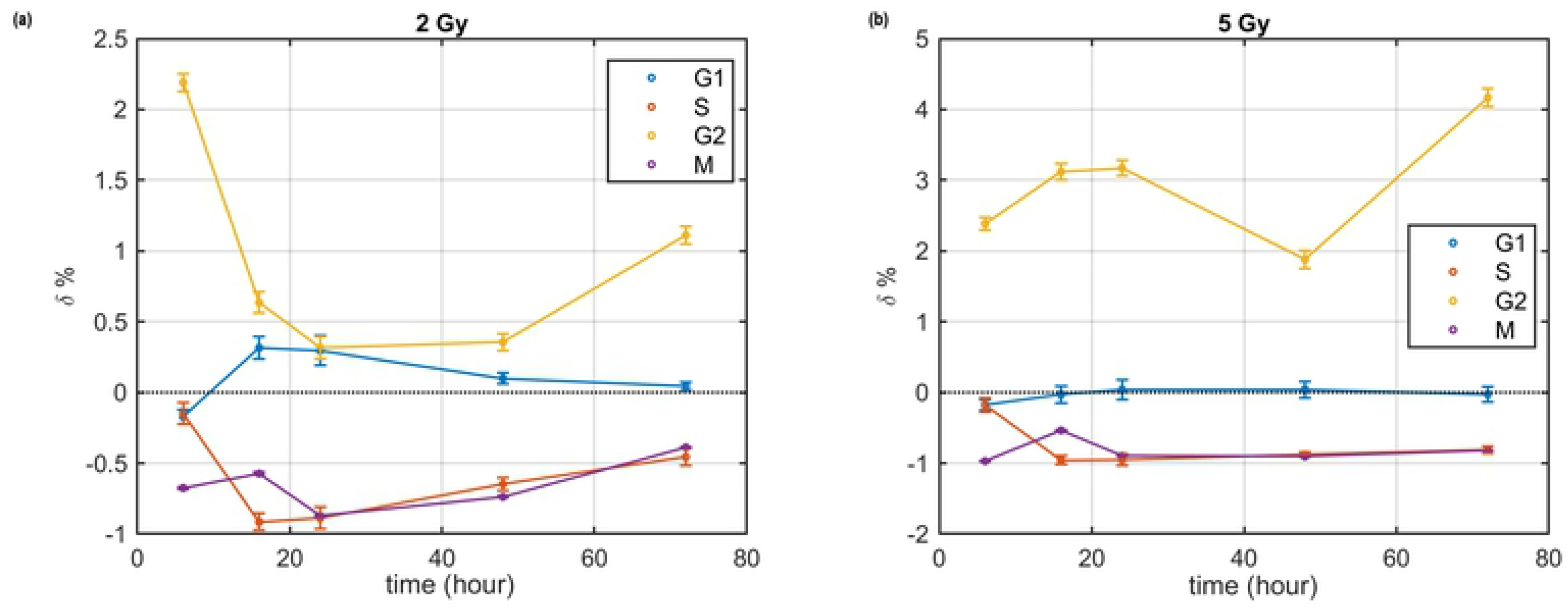
Relative differences between the doses and the sham condition. 2 Gy vs sham **(a)**, and 5 Gy vs sham **(b)**, as the percentages of cells in each phase as a function of time (lines are a guide for the eye).

From the figure we observe that the relative difference *δ* has a similar trend vs. time for the two doses as far as S, G_1_ and M-phases are concerned, though with different absolute values. The peak in G_2_-phase at 6 h for 2 Gy appears shifted to 16 h-24 h for the 5 Gy condition.

It has to be noted that relative differences with the sham condition at time points later than 24 h are affected by the fact that the unirradiated cells are approaching confluence, leading *e.g.* to the pronounced peak for the G_2_-phase at 72 h for 5 Gy.

### Data reproduction with the mathematical model

Starting from the work of Basse et al. 2003 [9], we extended a mathematical model for a population of cells describing their position within the cell cycle. The model is based on a system of partial differential equations that governs the kinetics of cell densities in all phases of the cell cycle, depending on time t and DNA content x and, only for DNA synthesis, on an age-like variable *τ*_*S*_ describing the time since arrival in S-phase. The explicit consideration of DNA content as a variable in the model allows, in principle, tuning of model parameter to reproduce experimental flow-cytometry profiles. In this section we briefly describe the general features of the model, we then turn to reproduction of experimental percentages of cells in cell-cycle phases, and extraction of model parameter values. Full details on the model formalism and on the strategy for data reproduction are given in the Materials and Methods section.

#### Mathematical model

The model is a classical compartmental model with four main compartments, one for each subpopulation of cells in a given cell-cycle phase. Four dependent variables: G_1_(x,t), S(x,t), G_2_(x,t) and M(x,t) represent the density number of cells with relative DNA content *x*, scaled to *x* = 1 in the G_1_-phase, at time *t*. The main parameters of the model are the transition rates that control the transfer of cells between phases. The dynamics between the compartments is represented by the schematic diagram in Fig. 5, and the model equations are the following (3):

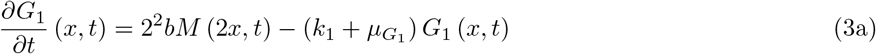

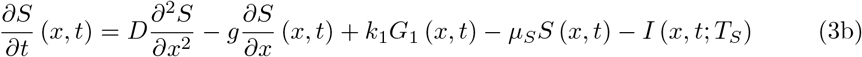

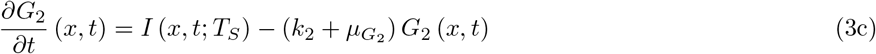

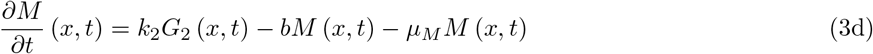

**Fig 5.**
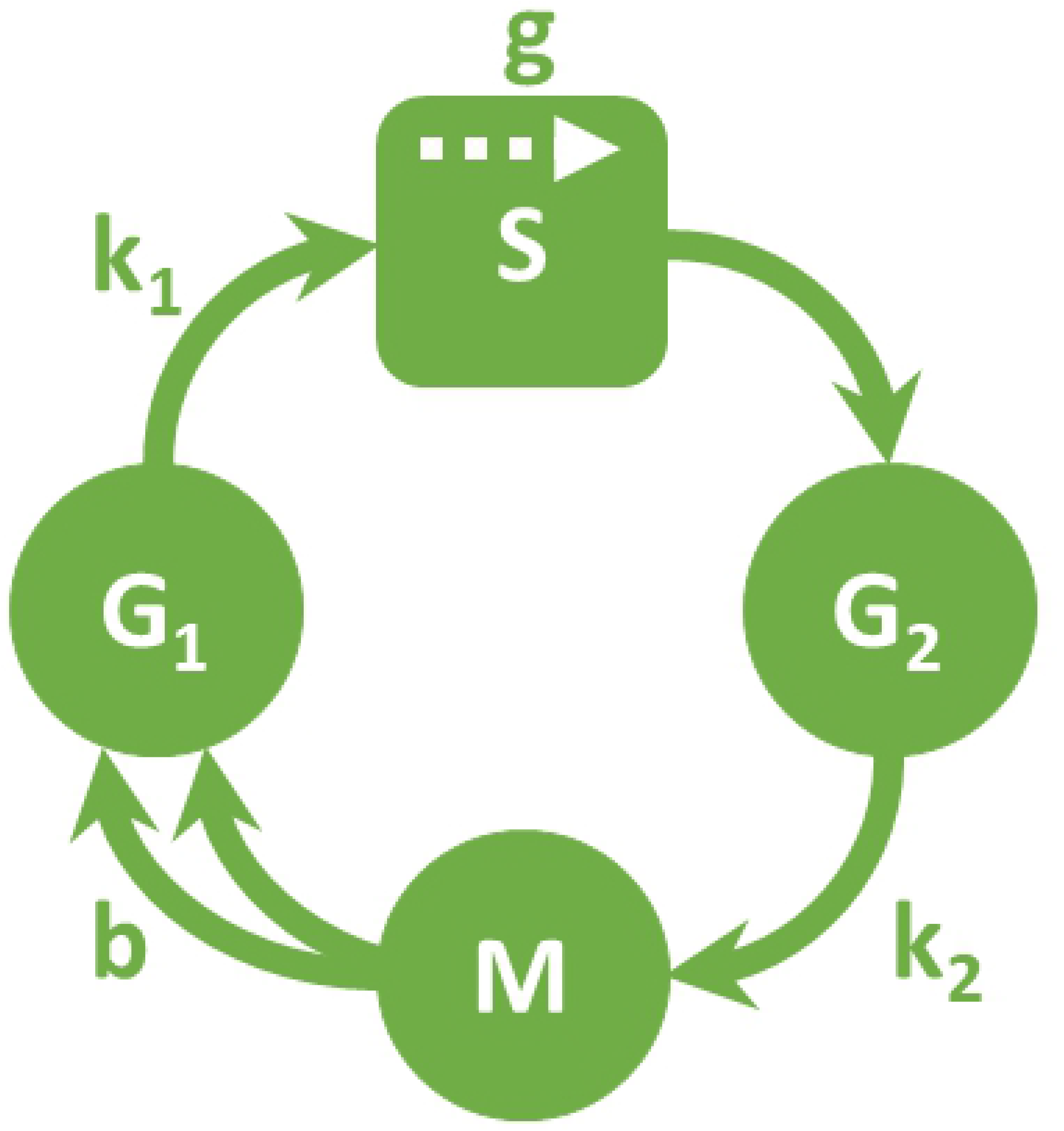
Schematic diagram of cell cycle model. *k*_*1*_ represents the transition rate from G_1_, *g* represents the DNA growth rate during S-phase,*k*_*2*_ represents the transistion rate from G_2_ to M-phase, *b* represents the division rate. Here as explained in the following section the *μ* parameters are set to 0, in order to describe vital cells only.

Each equation in (3) describes the variation of cell density in the corresponding phase. This is generally determined by a source term, that gives the influx of cells coming from the preceding phase, minus the loss of cells transitioning to the next phase and that due to cell death. The source term in G_1_-phase is given by the rate of divisions per unit time *b*. The S-phase requires a specific treatment: assuming that, on average, the time spent during DNA replication (T_S_) is constant for all cells, a cell entering the S-phase either exits after T_S_ hours or dies. In more detail, in addition to the influx from G_1_ and loss term due to death, in Eq. (3b) we have: the first term that is a dispersion term, taking into account the experimental variability of signal at increasing DNA content; the second term that describes the increase of DNA content at an average growth rate of *g* per unit time (instead, DNA content is constant in all other phases, so similar terms are not needed); the last term *I* (*x, t*; *T*_*S*_), that represents the subpopulation of cells which entered S-phase T_S_ hours before and are therefore ready to exit from S to enter G_2_. This latter term is the influx term in Eq. (3c) for the G_2_-phase. Full details on the model are given in the original work [9]. In normal conditions, cells are considered in a state of asynchronous balanced growth, where the percentage of cells in each of the four phases of the cycle remains constant. Experimentally, for an *in vitro* system, this corresponds to unperturbed exponential growth and requires cells to be far from confluence. This state represents the so-called steady DNA distribution (SDD) [10]. Rates of transition between phases are also constant. It has been shown that in such condition, the x variable is both uniquely determined by model parameters and independent of the initial distribution at time *t* = 0 [11]. A theoretical solution on the infinite temporal domain −∞ < *t* < ∞ exists, and asymptotically approaches the SDD condition [8]. For the purpose of this work, focusing on vital cells only, all μ_i_ parameters are set to 0. Also, a matricial formulation is adopted for the model implementation in MATLAB, as described in the Materials and Methods section.

#### Initial parameter values from experimental data

Experimental data on percentages of cells in cell-cycle phases for the sham condition are used to estimate model parameters for the SDD state. In Table 3 we report average percentages among the first 4 time points (including the 0 h pre-treatment sample), excluding time points for which cells get close to confluence (see Fig. 2). Using the experimental value of the doubling time T_D_, and an estimate of the time spent in mitosis as *T*_*M*_ = *T*_*D*_ · *f*_*M*_ (*f*_*M*_ is the average percentage of cells in M-phase), we can apply Steel’s formula (see Materials and Methods, Eqs. 17–20) and obtain initial guess values for model parameter, given in Table 4.

**Table 3.** Experimental percentages of cells in the 4 phases in the sham condition (cells in steady exponential growth)

| | $G_1$ | S | $G_2$ | M |
| --- | --- | --- | --- | --- |
| Experimental SDD | $61.06 \pm 1.59$ | $30.07 \pm 1.52$ | $8.07 \pm 1.01$ | $0.80 \pm 0.28$ |

**Table 4.** Values of model parameters with Steel formula’s (initial guess) and when fitted with *χ*^2^ minimization for the sham condition.

| | $k_1$ | g | $k_2$ | b |
| --- | --- | --- | --- | --- |
| Initial guess with Steel's formula | 0.0545 | 0.0813 | 0.2496 | 3.5798 |
| Fitted parameters | 0.0460 | 0.0829 | 0.2539 | 3 |

#### Model performance for the unperturbed condition

Given a set of initial parameters, the model can be run. For its evolution, a step-size in time h_t_ = 0.05 (h) is used. The model starts with 100% of cells in G_1_-phase. As a general criterion, we consider that convergence to a steady profile is reached when the sum of squared differences between percentages of all phases at time *t* and those at (*t* − 5) h becomes less than 10^−5^.

When running the model with initial parameters from Table 4, convergence is reached in T_SDD_ = 35.87 h. The corresponding model evolution and experimental percentages are shown in Fig. 6**-(a)**. In the SSD condition, model percentages are only close to experimental data, though within errors. Model parameters can therefore be adjusted, starting from initial guesses, to obtain a better reproduction of experimental data. This is done via *χ*^2^ minimization (see later in Materials and Methods, Eq. 21). Best fit parameters obtained via minimization are given in Table 4, and the evolution of the model and reproduction of the unperturbed conditions are shown in Fig. 6**-(b)**.

**Fig 6.**
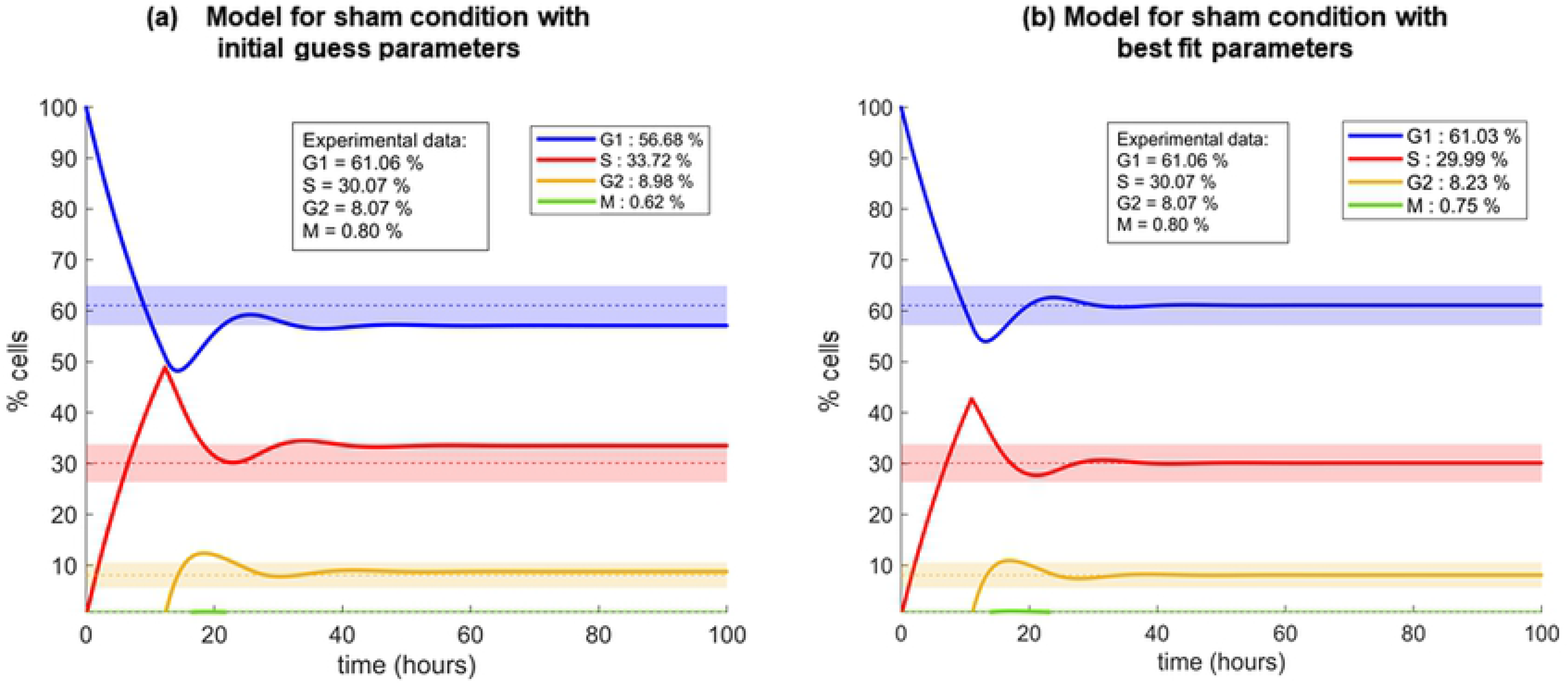
Model percentages of cells in each phase over time. Model percentages of cells in each phase with initial guess parameters by Steel (a), and with the optimized parameters (b). The dashed lines are the percentages of cells in each phase as measured experimentally (numerical values are also shown); the light coloured bands represent the experimental errors for the SDD condition.

#### Data reproduction strategy for the perturbed condition

The unperturbed cell population is growing exponentially at *t* = T_SSD_, before treatment. We then consider that cells are exposed to radiation at *t*′ > T_SSD_. For the sake of comparison to experimental data, we rescale the perturbation time *t*′ to 0. After the exposure, the effects of radiation are manifest as alteration of the transition rates between compartments.

A *χ*^2^ minimization strategy is not practical to find model parameters reproducing experimental data for the perturbed condition, as we justify in detail in Material and Methods section. Therefore, we use a simplified strategy for data reproduction, based on the following considerations:

1. alterations of *k*_*1*_ and *k*_*2*_, representing the transition probability from the gap phases to S or M, are expected to be more important. This is supported by biological evidence, as cell cycle checkpoints exist in G_1_ and G_2_-phases, and a larger impact on the population of such phases is observed in experimental data;
2. from the reproduction of data for the sham condition it is known that *g* and *b* parameters have less importance in the *χ*^2^ minimisation. We can assume that they are less modified by radiation, which is also supported by biological evidence, as S-phase and M-phase maintain an average constant duration.

To further simplify our strategy, we assume at first approximation that parameters variations can be described with step functions vs. time, thus altering their values in fixed time interval. We recall that the main features of the cell cycle perturbations that have been measured experimentally and that we want to reproduce are: an early G_2_ block (peaked at 6 h from irradiation for 2 Gy, and at a later 16 h time point for 5 Gy), and, after that, a strong increment of cells in G_1_-phase, that persists in time (see previous sections).

We find that the following alteration of model paramaters *k*_*1*_ and *k*_*2*_ lead to a good reproduction of experimental data:

1. for the 2 Gy condition, *k*_*2*_ is divided by a factor of 12 in the interval 0 h-6 h after irradiation. After that, from 6 h to 16 h, *k*_*2*_ is increased up to half its initial value, and later maintained constant throughout the simulation; at the 6 h time point *k*_*1*_ is divided by a factor 10, and maintained constant later;
2. for the 5 Gy condition, *k*_*2*_ is divided by a factor of 12 (as for 2 Gy condition) but this value is kept for a longer time, until 16 h after irradiation, to reproduce the longer block in G_2_-phase. At the 24 h time point, *k*_*2*_ is restored to 1/4 of its initial value, and later maintained constant. At 16 h, *k*_*1*_ is divided by 10 and later maintained constant.

Model calculations are shown in Fig. 7 and compared to experimental measurements. As it can be observed in the figures, simple changes in the parameters as a function of time lead already to a very good agreement between model and data. Note that, to provide a visual reference for phase percentages in the unperturbed condition, the time scale is shifted such that the irradiation time is at 10 hours in the plots.

**Fig 7.**
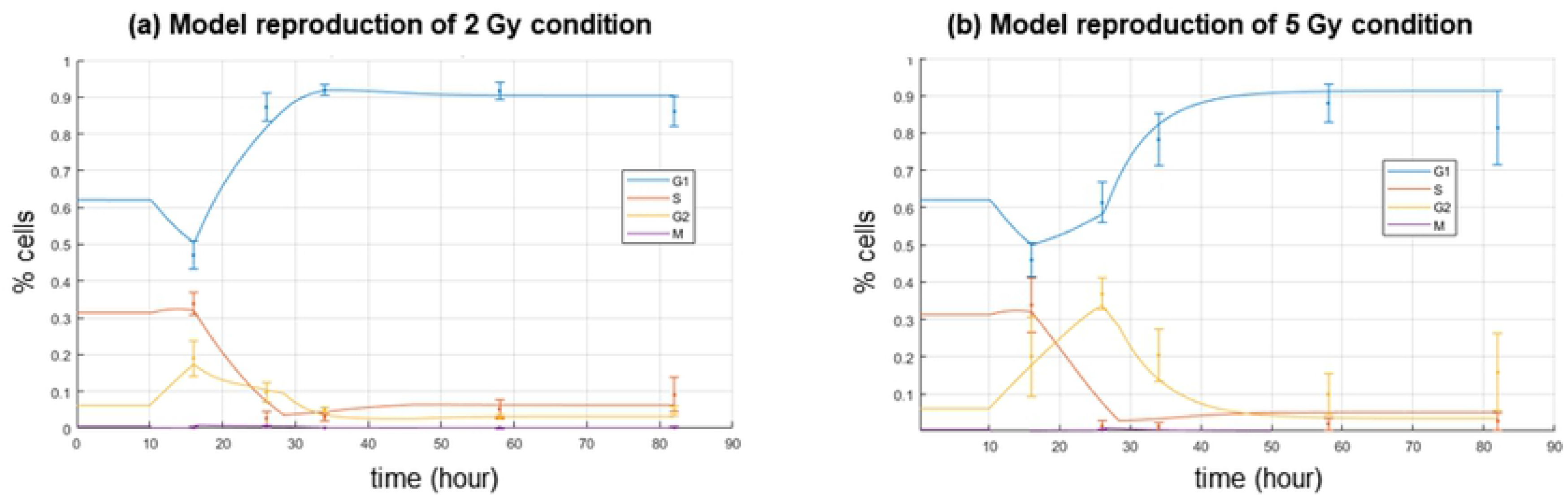
Model perturbation for cell cycle phases over time. Model perturbation for the 2 Gy condition **(a)**, and for the 5 Gy condition **(b)**. It is noted that the model is in the unperturbed condition up to 10 hours, at t=10 h cells are irradiated.

## Discussion

Starting from the work of Basse *et al.* [9], we implemented a computational model to reproduce how cells that survive radiation exposure are distributed in the cell cycle as a function of dose and time after exposure. To our knowledge, the original model had not been applied to cell cycle perturbations induced by ionizing radiation, but only tested for chemotherapeutic drugs.

This quantitative approach to model cell cycle perturbation underlies the following assumptions: transitions between phases are complete and irreversible [12]–[13]–[14]; and perturbations *e.g.* DNA damage, can affect the timing of these transistions. [15]. The model is deterministic, built with four compartments, each representing cells in a given phase of the cycle. The position in the cycle is given by the value of a DNA content variable, and the evolution in time is governed by a set of differential equations. Model parameters are transition rates (between G_1_- and S-phase and between G_2_- and M-phase), cell division rate and DNA synthesis rate. For the MATLAB implementation presented in this work we adopted a matrix formulation of the model, explicitly taking into account the time variable only, while information on the DNA content can be recovered *a posteriori* once the model is solved. The model gives as main output percentages of cells in each phase of the cycle, and a DNA content distribution can also be obtained, to be directly compared to cell-cycle profiles acquired with flow cytometry (namely, a distribution of signal intensity vs. a quantity proportional to DNA content).

The performance of the model was tested using data on IMR90 cells exposed to X-rays, with doses of 0 Gy (sham-irradiated or control), 2 Gy and 5 Gy, at multiple time-points after exposure: 6 h, 16 h, 24 h, 48 h and 72 h. Flow cytometry is used, obtaining information on cell percentages in each of the four phases in all investigated conditions. At difference with previous implementations of similar models, data include explicit discrimination of cells in G_2_- and M-phase, thanks to a flow-cytometry measurement protocol identifying M-phase through the detection of pH3 phosphorylation during mitosis.

Experimental results indicate that unperturbed cells in exponential growth settle over time to a steady DNA distribution, in agreement with expectations, in which the percentages of cells in each phase remain constant in time. Independent measurements of the doubling time (*i.e.* the overall time for a complete replicative cycle) provided a cross-check validation of the measured population increase rate. When cells are exposed to X-rays, a block in G_2_-phase in observed (at the early time point of 6 h after irradiation for 2 Gy and at 16 h for 5 Gy). After that, the percentage of cells in G_2_-phase decreases. The recovery to the pre-treatment condition is not total in both cases and seems to be dose dependent. At the same time cells accumulate in G_1_-phase right after the block in G_2_-phase, with a modification that seems persistent in time.

Model parameters can be adjusted to reproduce experimental data. As a general strategy, we obtained the set of model parameters needed to reproduce the sham condition, and assume that any perturbation can be described by the alteration of these parameters. The trends observed in experimental data are reproduced by varying only two of the parameters: *k*_*1*_ and *k*_*2*_, respectively the transition rate from G_1_- to S-phase and from G_2_- to M-phase. This is consistent with the biological knowledge of the main role of the two checkpoints in the gap phases to control the cell cycle progression. Interestingly, as the analysis is limited to living cells, the model parameters (giving the entity of the perturbation) are not so different in the two perturbed conditions. We observed instead a difference on the duration of the blocks in gap phases, and on the ability to recover to the unperturbed condition. Therefore, a perturbation of similar entity was applied for both doses, but kept for longer times for the higher dose, with a later recovery to a lower value at 5 Gy compared to the 2 Gy. Similarly, the same perturbation is applied to *k*_*1*_ but a later time point (16 h vs. 6 h) for the highest dose condition, probably representing (more than a block in G_1_-phase) the renewed influx to G_1_ of cells surviving the G_2_ block, which coherently happens later for the higher dose.

How parameters are varied in order to reproduce data also gives an insight on possible biological interpretations: up until 4 - 6 h post irradiation only a slowing of S-phase entry is observed, even at high doses (10 Gy), and this can be explained by the principal molecular mechanisms for G_1_ arrest, that is a slow process involving transcriptional activation by p53 of p21 that leads to inhibition of pRb phosphorylation and G_1_ arrest [16–19].

In this work a *χ*^2^ fit strategy involving all the experimental time-points is used for the reproduction of the unperturbed condition only, while for the irradiated conditions a minimization is performed in discrete time intervals, giving as results step functions for parameter variation as a function of time. Perturbed parameters therefore represent average rates in discrete time interval. A first improvement of the reproduction strategy of the perturbed data would be to refine the time dependence of the perturbed parameters, introducing linear functions with imposed continuity conditions between time points. Furthermore, it would be necessary to implement a *χ*^2^ fit strategy also for the perturbed conditions that could include all time-points simultaneously. This deserves further investigation as it is expected that the objective functions will have a complex behaviour as a function of model parameters and that the perturbed biological system is less easy to characterize with additional constraints (as the measurement of the population increase rate). Optimization of the reproduction of the shape of the cell cycle profile could also be performed, fully exploiting the flow-cytometry data. In this simulation it is assumed that the variances increase linearly from 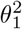 to 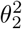 during S-phase, doubling in value, with a coefficient of variation of 5%, compatible with the experimental CV obtained by flow-cytometry.

It has to be stressed that results presented in this work in terms of percentages of cells in the different phases of the cycle always correspond to the fraction of living cells only. The importance of cell death cannot be quantified from our dataset. This information could be added experimentally and used as input for further model developments: the model could be expanded introducing a new compartment of dying cells. This same formulation of the model could be already adapted to reproduce cell death data adjusting the values of death rate parameters (currently set to 0). Dedicated experimental flow-cytometry protocols must be planned, *e.g.* using Annexin V with FITC vs. Propidium Iodide, that can discriminate cells based on viability, and possibly identify the death mechanism (apoptosis vs. necrosis). Additionally, a quiescent cell compartment could be added to the model to take into account the existence of the G_0_-phase. What we observe in our dataset as an increase in the population of G_1_ after irradiation could also be due to cells that exit the replicative cycle and accumulate in a quiescent G_0_-state.

To conclude, the computational model we present in this work has the potential to reproduce perturbations of cell cycle following exposure to ionizing radiation, as well as to possible other agents as drugs. We here focus on the percentage of living cells in different phases of the cycle as a function of radiation dose and time, offering a proof-of-concept validation of the model with flow-cytometry experimental data on a chosen cell line. The model can be easily applied to reproduce different datasets, varying cell line, radiation dose, time points, changing the type of radiation or even testing different agents, as mentioned. Biological insight on underlying response phenomena can be gained, comparing model parameters that are necessary to reproduce different datasets. By construction, the model can be easily extended to the consideration of cells exiting the cycle, either undergoing cell death or transition to a quiescent state. New dedicated datasets should be acquired to benchmark possible model extensions. A model of this kind constitutes the core of more complex tools for pre-clinical/clinical use, that are able to include cell cycle perturbation for predictions in radiotherapy, *e.g.* giving information on optimal fractionation schemes or on the possible synergistic combination of radiation and chemotherapeutic drugs.

## Materials and methods

### Experimental cell cycle analysis

#### Cell culture

In this work a primary culture of IMR90 human fibroblast cells (ATCC^©^ CCL186^®^, USA) originated from lung tissue is used. Cells were cultured in complete medium (EMEM supplemented with 12.5% of FBS, 1% of L-glutamine 2 mM and 1%of NEAA) at 37 °C and 5% CO_2_, until confluence was reached at 80-90%. Cells used for the measurements were always between the 8^th^ and the 25^th^ passage. IMR90 cells were plated at a density of 10^5^ per T25 flask (GreinerBio-One, Germany).

#### Growth Curve

Indipendent measurements on cellular growth curves were performed in order to estimate the doubling time (T_D_) of an unirradiated population. 10^5^ cells were seeded per T25 flask, they were cultivated and harvested at following time points: 6 h, 16 h, 24 h, 48 h, 72 h (the same time points for the following study of the radiation effects). The samples were trypsinized, 10 μl of the pellet was placed in a Bürker counting chamber. The number of cells were counted under an inverted microscope. Experiments were repeated in biological triplicate, each of them was performed in technical duplicate. The data were fitted to the following exponential fucntion via non-linear least squares methods: *fit*(*t*) = *N*_0_ exp(*γt*), where *N*_*0*_ is the number of cells at *t* = 0, and *γ* is the growth rate.

#### Irradiation setup

Irradiations were performed at the Radiotherapy facility of IRCCS Istituti Clinici Maugeri (Pavia, Italy), with a 6 MV linear accelerator Clinac (Varian, USA), regularly used for radiotherapy treatments. Cell samples were exposed to X-rays at two different doses of 2 Gy and 5 Gy, with a dose rate of 3 Gy/min. Irradiations were performed as previously described in detail [20]. Control samples, so-called “sham” samples, underwent the same environmental stress as the irradiated samples, except for exposure to radiation. After irradiation samples were transferred in the incubator at 37 °C with 5% CO_2_ level. At each time point (6 h, 16 h, 24 h, 48 h, 72 h) cells were treated for flow-cytometry measurements, as described in the next section. Experiments were repeated in biological triplicate, each of them was performed in technical duplicate.

#### Cell cycle characterization

After irradiation, cells were incubated with 2 *μg/μl* EdU (Click-i™ Plus EdU Alexa Fluor™ 488 Flow Cytometry Assay Kit, Invitrogen, USA) for 1 h, then fixed following the kit manufacture instructions with minor changes, to discriminate S-phase. M-phase was discriminated with the phospho-Histone H3 (Ser-10) primary antibody (1:25 in PBS with 1% of NGS, Cell Signaling Technology, USA [RRID:AB 331748]) and the secondary anti-mouse IgG antibody (1:500, Alexa Fluor 555 Conjugate, Cell Signaling Technology, USA [RRID:AB 1904022]). FxCycle Violet dye (4’,6-diamidino-2-phenylindole, dihydrochloride) was used to measure the total DNA content, following manufacture instructions. The Attune NxT Acoustic Focusing flow cytometer (Thermo Fisher Scientific, USA) employed for these experiments is equipped with two lasers emitting, respectively, in the blue (488 nm, 50 mW) and in the violet (405 nm, 50 mW).

The full gating strategy to discriminate the four cell cycle phases is as follows: after the identification of the cell population and of singlets (Fig. 8, panel (a) and (b), respectively), the characteristic cell cycle spectrum, as measured with FxCycle™ Violet, (Fig. 8, panel (c)) is analyzed. A further gate is applied, to select the whole spectrum where the signal of FxCycle™ Violet is proportional to the amount of DNA: this has the typical form of two Gaussians, the second with a mean value equal to the double of the first peak, and a plateau in between. These three distributions represent respectively cells in G_1_-phase, G_2_/M-phase and S-phase.

**Fig 8.**
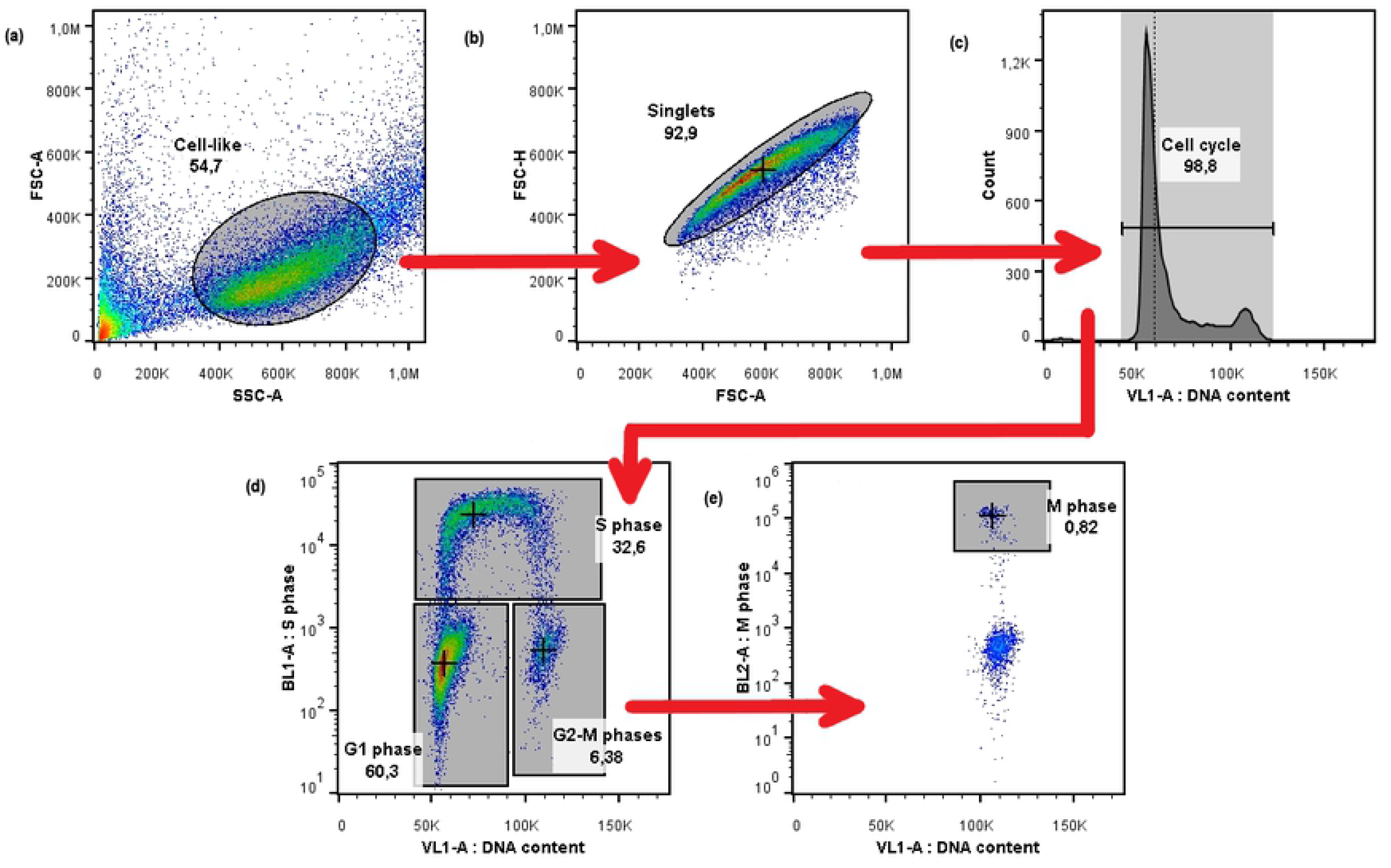
Flow cytometry gating strategy. Gating hierarchy of a representative sample (Sham 16 h). From left to right and from up to down: **(a)** in FSC-A vs SSC-A, identification of the cloud of cell-like events; **(b)** in FSC-A vs FSC-H, identification of singlets cells; **(c)** histogram of VL1-A shows the cell cycle profile; **(d)** in VL1-A vs BL1-A, three distinct groups (cells in G_1_-phase, S-phase, and G_2_/M-phases) are shown, and finally from the last gate in **(e)** it is shown the separation of M-phase from the rest of the cells in the VL1-A vs BL2-A plane.

The cell cycle spectrum in VL1-A channel alone cannot provide a quantitative discrimination of the four phases, so stainings with EdU and pH3 antobody were introduced. In the bi-parametric plane BL1-A vs. VL1-A (EdU vs. FxCycle™ Violet, Fig. 8 panel (d)) there are three distinct clouds of events: the cloud in the bottom left represents cells in G_1_-phase, that have a low signal of FxCycle™ Violet and low signal of EdU. The events in the bottom right part of the graph represent cells in G_2_ or M-phase, while the horseshoe-like subset with a high signal of EdU represents cells in S-phase. The events collected in the gate “G_2_-M”are visualized in the VL1-A vs. BL2-A plane (pH3 vs. FxCycle™ Violet): here events with a double positive fluorescence correspond to cells in M-phase, that have a double content of DNA and a high content of phosphorylated H3 (Fig. 8 panel (e)).

#### Statistical analysis

Data acquisition was performed with the Attune NxT software. Data analysis was carried out with FlowJo^®^, a specialised software used for flow-cytometry applications. As a result of the gating procedure described above, we obtained cell numbers in each of the four cell cycle phases. Data were then expressed in terms of percentages with respect to the whole cell population (hence normalized to the sum of cells in all gates used to identify cell cycle phases). Data are given as average between both biological (3) and technical (2) replicates, and all experimental uncertainties reported are intended at 1*σ*.

### Mathematical model details

#### Mathematical model: boundary and initial conditions and steady-state solution in unperturbed condition

Model equations (3) govern the dynamics between cell-cycle phases. To solve the system and find the time *t* and DNA content *x* dependence of the four phases, G_1_(x,t), S(x,t), G_2_(x,t) and M(x,t), boundary and initial conditions are needed, that we now discuss.

The following boundary condition (Eq. (4)):

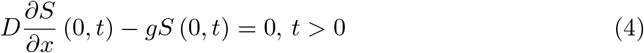

ensures a positive DNA content in all cells at all times.

At first, as initial conditions, all cells are synchronised in G_1_-phase, while the other three compartments are empty. An approximation of the flow-cytometric profile of G_1_-phase at time t is given by Eq. (5):

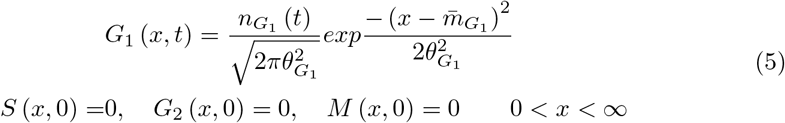

where 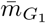 is the mean DNA content in G_1_-phase and 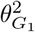 is the corresponding variance. For simplicity, the mean parameter is normalized to a relative value *x* = 1 for G_1_-phase, thus giving *x* = 2 for G_2_-phase and M-phase, while the variance is chosen sufficiently small so that *G*_1_ (*x*, 0) exists only for *x >* 0, and can be adapted to simulate the experimental variance of the flow-cytometric profile.

Given the initial conditions as Eq. (5), the model is made evolve until T_SDD_ hours to reach a steady DNA distribution. After such time, the model is considered to give the cell cycle distribution of cells in exponential growth. The variance 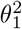 for the Gaussian distribution of cells in G_1_-phase is fixed at 0.05, which is chosen based on the experimental variance of the G_1_-phase sham profiles (around 5% of the mean). The variance in G_2_-phase and M, 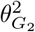, is considered as two times 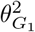. Starting from the initial condition, Fig. 9 shows the cell cycle profile evolution in time. In practice, the full profile in *x* is superimposed to the solution of the problem in its matricial formulation, that gives the number of cells in each phase at each time (see next section).

**Fig 9.**
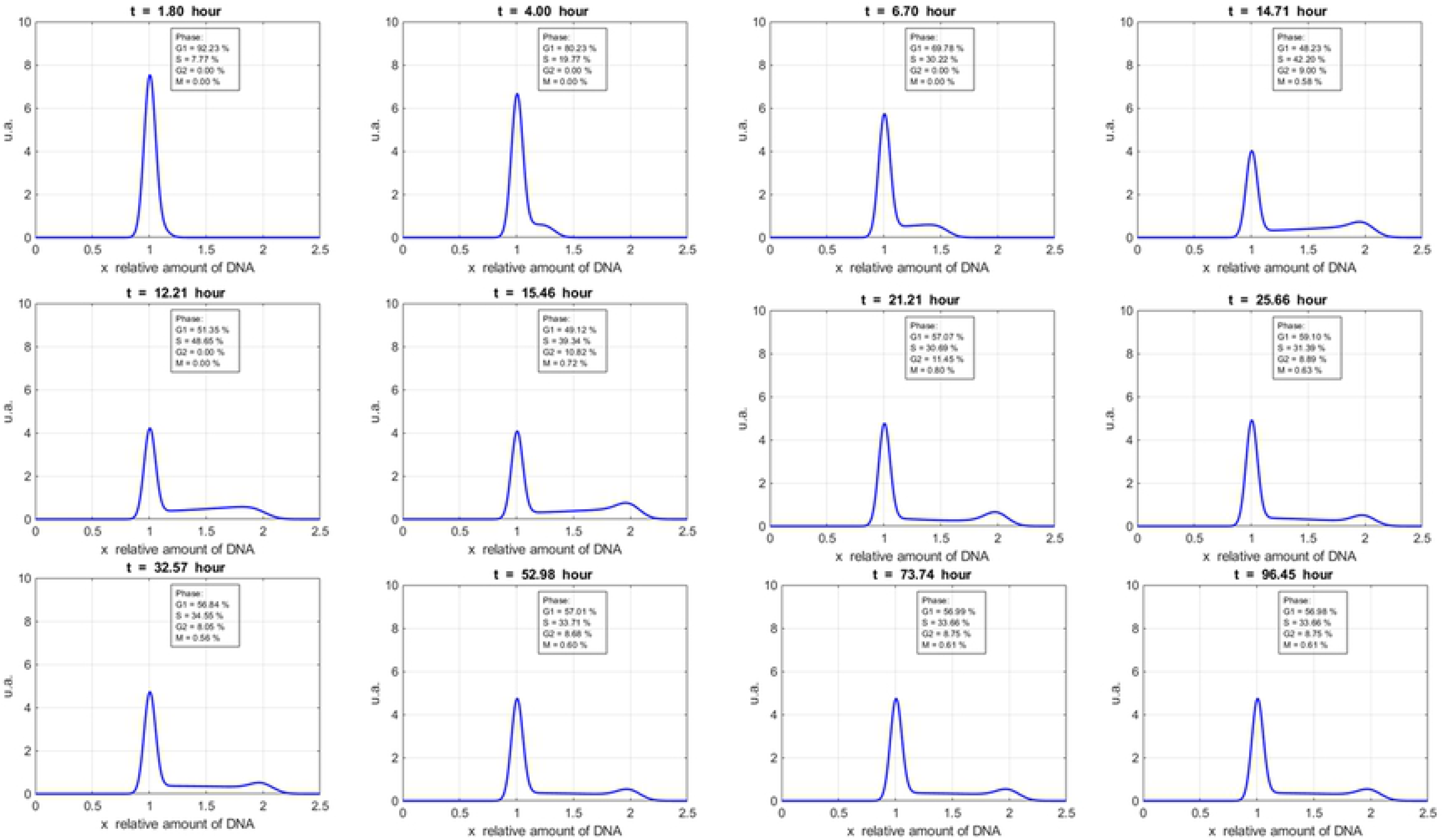
Model evolution in time. Snapshots in time of the model evolution of cell cycle profile along x (DNA content).

#### Matricial model formulation

For a practical implementation of the model in MATLAB we resort to a matricial model formulation, integrating Eq.s (3) over the DNA content domain from 0 to X, with X upper limit for the *x*-axis, large enough to simulate infinity so that the difference of the integrations over 2X and over X is negligible. The matrix system is resolved using finite forward differences method, with discretization in the time variable. The integration leads to the following system of temporal ordinary and partial differential equations:

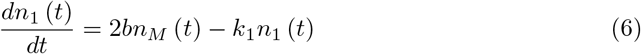

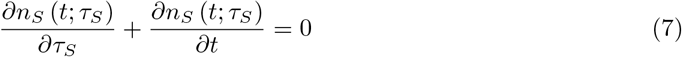

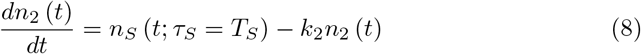

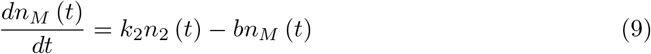

where:

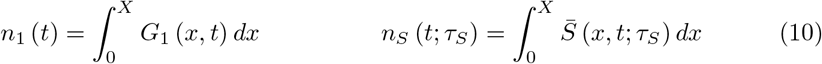

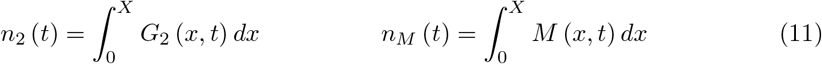

Then, Eqs. 6–9 are rewritten as a linear system of equations and finite forward differences method is applied to resolve it:

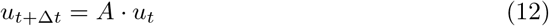

with:

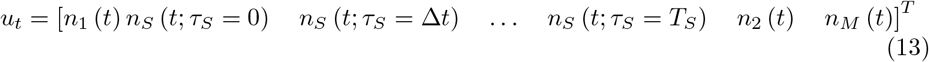

and

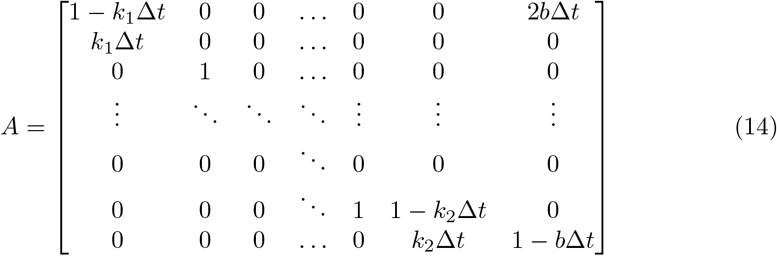

The eigenvector corresponding to the eigenvalue of largest magnitude of the transition matrix A gives the long-term distribution of cells. The eigenvalue determines the rate of population growth and allows to compute the doubling time of the cell population [21]. In the discretized matrix form of the model of an unperturbed cell line, the number of cells in each phase can be tracked over time without any knowledge of the DNA distribution in each phase. This formulation is more convenient for practical model implementation, but it provides no information on the flow cytometric profile to be compared to data. To obtain this information at any time in each phase, normal distributions can be superimposed to data from the matrix model. It is assumed that experimental flow cytometric profiles obtained from a homogenous cell population are approximately Gaussian, and experimental variances can be used to adapt theoretical distributions. For G_1_-phase at the arbitrary time *t*′, the flow cytometric profile is:

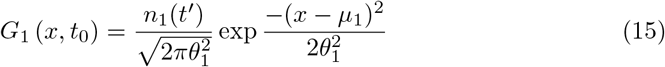

The mean μ_1_ for G_1_-phase is chosen at a relative DNA content *x* = 1. The S-phase profile is composed by a sum of Gaussian distributions with mean values shifted along the *x*-axis, proportional to *τ*_*S*_ (the time each cell has spent in S-phase). So, the mean in S-phase is μ_1_+g_S_*τ*_*S*_. For G_2_ and M, the profile is a Gaussian function like the one in Eq. 15, but with mean respectively μ_2_ and μ_M_ at *x* = 2 [22]. It is assumed also that the variances increase linearly from 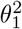 to 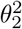 during S-phase. As mentioned, the main motivation to consider this discretized formulation of the model is the faster computational time when using a language as MATLAB.

#### Initial parameter values from experimental data

For a first estimate of model parameter values in the unperturbed condition we can start from the so-called Steel’s formulas [23]: such formulas give the average time spent in each phase (T_phase_) as a function of measured percentages of cells in cell-cycle phases (*f*_*phase*_) and the doubling time T_D_. Formulas are as follows (Eq. 16):

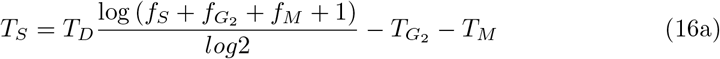

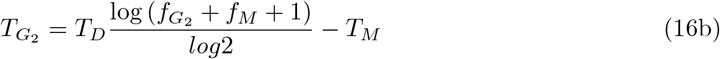

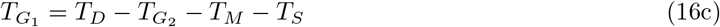

where T_M_ can be separately estimated as T_D_ · *f*_*M*_.

As previously mentioned, for a population of cells with a steady DNA distribution, *f*_*phase*_ percentage values are unaltered in time. Steel’s formulas are then combined with the relation between the average time in each phase and the model parameters as in [24]:

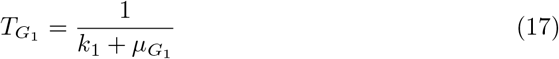

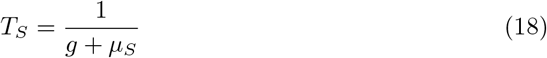

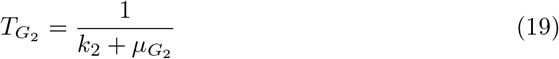

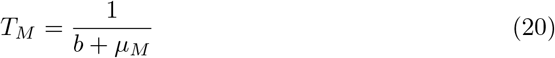

If μ_i_ parameters determining cell death in Eq.(3) are set to 0, transition rates are simply the reciprocal of the average time spent in each phase: if a block occurs in a particular phase, the time spent in that phase will increase and the rate transition will tend to zero.

#### Fit strategy for the unperturbed condition

Parameter values obtained from Eqs.17–20 are used as an initial guess. A minimization strategy is then necessary, to obtain the set of parameters that allow the best reproduction of experimental data for the unperturbed condition. The minimization strategy is not uniquely defined, and possible objective functions show multiple local minima, with different sets of parameters returning similar profiles, with very different doubling time.

We follow a sequential minimisation procedure on the four parameters: *k*_*1*_, *g*, *k*_*2*_, *b*. We define the objective function to minimize the difference between experimental and theoretical percentages as:

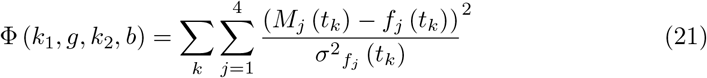

where [k_1_, g, k_2_, b] is the vector of the model parameters spanning between their limits, *M*_*j*_(*t*_*k*_) is the theoretical percentage of cells in the phase *j* at time *t*_*k*_, *f*_*j*_(*t*_*k*_) is the experimental percentage of cells in the phase *j* at time *t*_*k*_ and 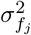 is the corresponding experimental error.

The following constraints are imposed: *k*_1_ ∈ [0, 1], *g* ∈ [0.05, 0.5], *k*_2_ ∈ [0, 1] and *b* ∈ [0, ∞). These constraints ensure positivity of the parameter values, reflect the fact that *k*_*1*_ and *k*_*2*_ are probabilities and assume that the time spent in S-phase is approximately between 2 and 20 hours [25]. Lower and upper bounds of the fit parameters kept during minimization are given in Table 5.

**Table 5.** Bounds of the four parameters to be fitted.

| Parameter | [dim] | Lower bound | Upper bound |
| --- | --- | --- | --- |
| $k_1$ | $\left[\frac{1}{t}\right]$ | 0.03 | 0.06 |
| $g$ | $\left[\frac{x}{t}\right]$ | 0.05 | 0.5 |
| $k_2$ | $\left[\frac{1}{t}\right]$ | 0.05 | 0.5 |
| $b$ | $\left[\frac{1}{t}\right]$ | 3 | 5 |

It has to be recalled that the four parameters are related: the sum of their reciprocal gives the overall duration of the cell cycle, hence what is experimentally measured as the doubling time. Therefore, this can be used as a criterion to drive the optimization: any local minimum of the objective function for which the derived cell cycle length is far from our estimation can be discarded. The same holds for minima that occur for parameter sets too far from initial conditions, as initial parameter guesses provided by Steel’s formula have been shown to give a reasonable agreement with experimental data. For all these reasons, the minimization is performed varying parameters in relatively narrow intervals, in particular for *k*_*1*_, as the objective function is found to have a strong dependence on its value. On the contrary, minimization is less affected by the values of *b* and *g*. Taking also into account the lower importance of the *b* parameter and the low values assumed by M-phase percentages, in a first instance the parameter was fixed to the value 3.

The optimisation of the fit (minimization of Φ) for the unperturbed condition is performed with a sequential quadratic programming method via the MATLAB routine fmincon [26].

## Acknowledgments

The authors acknowledge: Dr. Jacopo Morini and Dr. Gabriele Babini for initial discussion and suggestions on the experimental design; Dr. Marco Liotta and Dr. Paola Tabarelli de Fatis for performing the irradiation at *Fondazione IRCCS Maugeri*, Pavia, Italy.

